# Retinal and cortical visual acuity in a common inbred albino mouse

**DOI:** 10.1101/2020.11.03.366385

**Authors:** Michelle Braha, Vittorio Porciatti, Tsung-Han Chou

**Author notes:** Corresponding author (THC).

## Abstract

While albino mice are widely used in research which includes the use of visually guided behavioral tests, information on their visual capability is scarce. We compared the spatial resolution (acuity) of albino mice (BALB/c) with that of pigmented mice (C57BL/6J). We used a high-throughput pattern electroretinogram (PERG) and pattern visual evoked potential (PVEP) method for objective assessment of retinal and cortical acuity, as well as optomotor head-tracking response/ reflex (OMR). We found that PERG, PVEP, and OMR acuities of C57BL/6J mice were all in the range of 0.5-0.6 cycles/degree (cyc/deg). BALB/c mice had PERG and PVEP acuities in the range of 0.1-0.2 cyc/deg but were unresponsive to OMR stimulus. Results indicate that retinal and cortical acuity can be reliably determined with electrophysiological methods in BALB/c mice, although PERG/PVEP acuities are lower than those of C57BL/6J mice. The reduced acuity of BALB/c mice appears to be primarily determined at retinal level.

## Introduction

The study of visual capabilities in inbred mouse strains is of growing importance, as many behavioral tests used to characterize the effects of genetic manipulations rely on visual information [1]. BALB/c mice are among the most widely used inbred strains in animal experimentation, including investigating visual system abnormalities in autism [2]. Vision in albino mice has long been subject of interest since oculocutaneous albinism in all mammalian species is associated with poor vision [3] due to developmental abnormality in ocular melanin synthesis, resulting in retinal underdevelopment and misrouting of the visual pathway [4]. Few studies have investigated vision capabilities in albino mice, however with some inconsistency. Some studies using virtual-reality optomotor head-tracking response/ reflex (OMR) in BALB/c mice were unable to elicit a measurable response [5, 6], while in another study in BALB/c mice [7], investigators were able to measure a small OMR in the direction opposite to that of the visual stimulus, resulting in a visual acuity (0.12 cyc/deg) lower than that reported for pigmented C57BL/6 mice (B6) (0.5-0.6 cyc/deg)[8]. BALB/c mice, in contrast to other common inbred strains, are reported to be unable to learn a two-alternative swim task (VWT) to evaluate visual acuity [9], while another study used the VWT to assess the visual acuity (0.3 cyc/deg) of BALB/c mice [7].

The Pattern Electroretinogram (PERG) [10] and pattern visual evoked potentials (PVEPs) have been extensively used to measure acuity and contrast thresholds in both wild-type and mutant mice [11-13]. In the present study we have used PERG and PVEP to assess visual acuity at retinal and cortical level in BALB/c and B6 mice and have also compared PERG/PVEP results with corresponding assessments obtained with OMR.

## Methods

### Visual acuity assessment by pattern electroretinogram (PERG)

The study was performed in B6 mice (4 months old, n=10, 20 eyes) and BALB/c mice (4 months old, n=10, 20 eyes) purchased from Jackson Labs (Bar Harbor, ME, USA). All procedures were performed in compliance with the Association for Research in Vision and Ophthalmology (ARVO) statement for use of animals in ophthalmic and vision research. The experimental protocol was approved by the Animal Care and Use Committee of the University of Miami (Project protocol number: 16-247). All mice were maintained in a cyclic light environment (12 h light: 50 lux–12 h: dark) and fed with a grain-based diet (Lab Diet, 500, Opti-diet; PMI Nutrition International, Inc., Brentwood, MO, USA). The PERG was recorded simultaneously from each eye using one common subcutaneous stainless steel needle in the snout as previously described [14]. In brief, mice were anesthetized by means of intraperitoneal injections (0.5–0.7 mL/kg) of a mixture of ketamine (42.8 mg/mL) and xylazine (8.6 mg/mL). Mice were gently restrained using a mouth bite bar and a nose holder that allows unobstructed vision and kept at a constant body temperature of 37.0°C using a feedback-controlled heating pad. Under these conditions the eyes of mice are naturally wide open and in a stable position, with pupils pointing laterally and upward. Visual stimuli were black-white horizontal gratings of 100% contrast and of different spatial frequencies (0.047 – 0.571 cyc/deg in eight steps) generated on two (15 cm x 15 cm) LED displays (mean luminance 700 candela/m^2^ (cd/m^2^) for visual acuity testing at high luminance or 70 cd/m^2^ for visual acuity testing at low luminance), and presented to each eye independently (Jorvec Corp, Miami, FL, USA). Pattern stimuli reversed at slight different rates for each eye (OD, 0.984 Hz; OS, 0.992 Hz) to allow mathematical deconvolution of the response and isolation of independent monocular PERG using one channel continuous acquisition and phase-locking average [14]. For each spatial frequency, the PERG and noise amplitudes were automatically measured from peak to trough in the time window 50-350 ms. PERG amplitudes for all mice were plotted as a function of spatial frequency and data interpolated with a linear regression. The spatial frequency at which the linear regression of PERG amplitude intercepted the median noise level was considered as the PERG spatial resolution (PERG acuity). For each mouse, the complete recording sequence (lowest to highest spatial frequency) required about 40 minutes. Representative examples of PERG waveforms and noise waveforms in B6 and BALB/c at different spatial frequencies are shown in fig 1A. In order to have an estimate of PERG acuity that was not based on the assumption of linearity between decreasing PERG amplitude and increasing spatial frequency, we calculated for each SF the signal-to-noise ratio (SNR). SNRs > 1 were assigned a categorical value of YES while SNRs ≤ 1 were assigned the value of NO. Data were then submitted to a logistic analysis to retrieve the spatial frequency at which YES responses had 50% probability.

**Fig 1.**
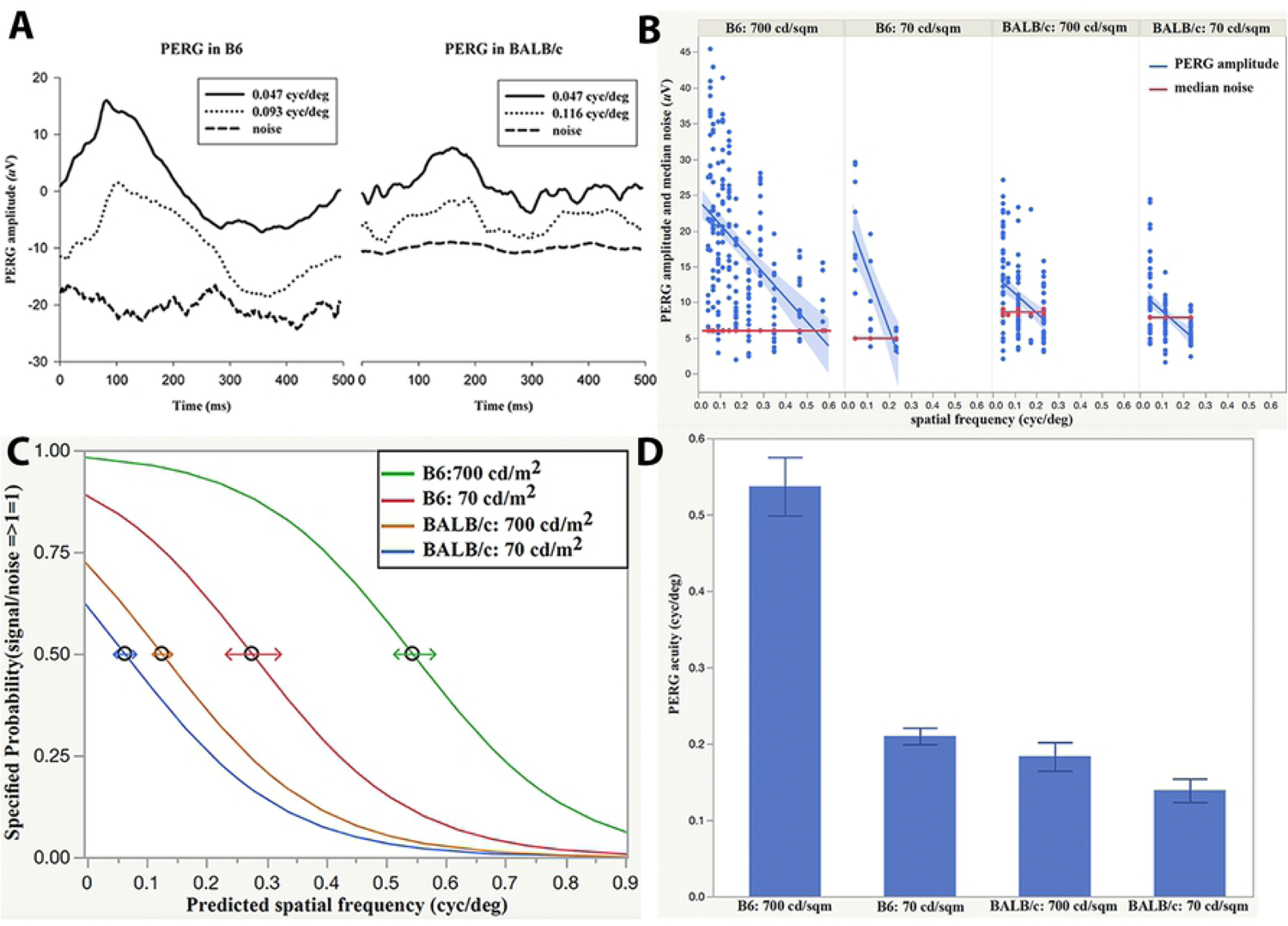
Comparison between PERG acuity of C57BL/6J and BALB/c mice. (A) Examples of PERG and noise waveforms from B6 and BALB/c mice at different spatial frequencies and 700 cd/m^2^ mean luminance. (B) PERG amplitudes as a function of spatial frequency for all mice. Linear regressions of PERG amplitude (blue lines with 95% confidence region) intersect the median noise level (red lines) at different spatial frequencies depending on strain and mean luminance. (C) Group PERG acuity assessed with categorical logistic method (50% chance of PERG/noise ratio >1; horizontal arrows represent 50% confidence interval) showing that acuity depends on strain and luminance. (D) Mean PERG acuity (± SEM) in B6 and BALB/c calculated by linear extrapolation of PERG amplitude to the median noise in the individual mice.

### Visual acuity assessment by pattern visual evoked potential (PVEP)

The study was performed in a subset of B6 mice (4 months old n=5, 10 eyes) and BALB/c mice (4 months old, n=5, 10 eyes) purchased from Jackson Labs (Bar Harbor, ME, USA). PVEPs were recorded in anesthetized mice as described in PERG methods section, with the difference that recording electrodes were stainless steel screws (shaft length 2.4 mm, shaft diameter 1.57 mm; PlasticsOne, Roanoke, VA, USA) inserted into the skull, contralateral to the stimulated eye 2 mm lateral to the lambda suture, which corresponds to the monocular visual cortex [11]. Reference electrodes were also stainless steel screws inserted into the skull to the ipsilateral to the stimulated eye 2 mm lateral to the lambda suture. The ground electrodes were subcutaneous stainless steel needles at the base of the tail. PVEP signals were amplified (10,000) band pass filtered (1–100 Hz) and averaged (two consecutive, partial averages of 600 epochs each, whose grand-average constituted the PVEP response) [15]. For each spatial frequency, PVEP and noise amplitudes were measured from peak to trough. PVEP amplitudes for all mice were plotted as a function of spatial frequency and data interpolated with a linear regression. The spatial frequency at which the linear regression of PVEP amplitude intercepted the median noise level was considered the PVEP spatial resolution (PVEP acuity). A representative example of PVEP waveforms and noise waveforms at different spatial frequencies is shown in fig 2A. As for the PERG, PVEP acuities were also calculated using logistic regression of categorical variables (SNR >1: YES; SNR ≤ 1: NO) as a function of spatial frequency. PVEP acuity was considered 50% probability of YES responses.

**Fig 2.**
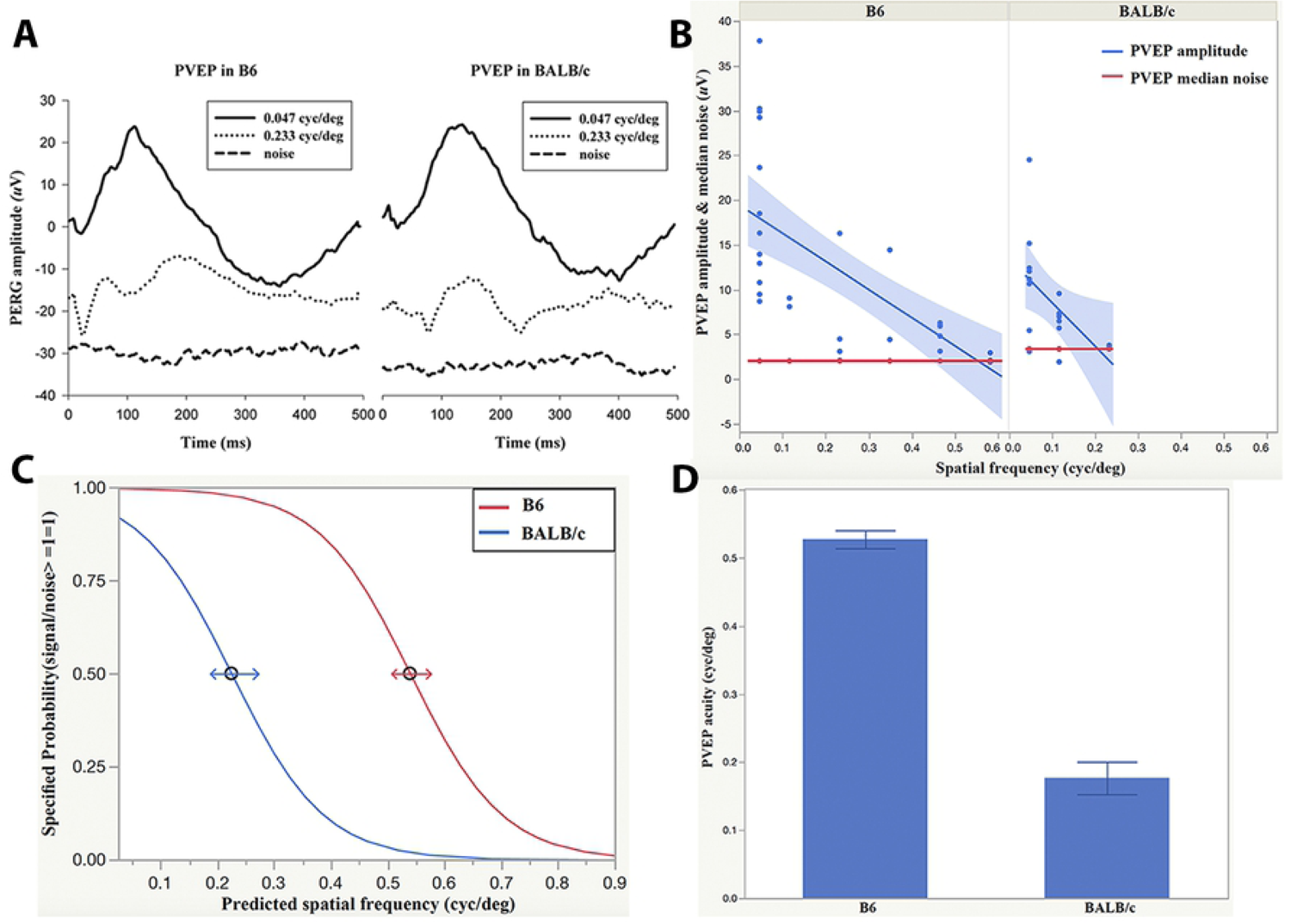
Comparison between PVEP acuity of C57BL/6J and BALB/c mice. (A) Examples of PVEP and noise waveforms recorded from B6 and BALB/c mice at different spatial frequencies and 700 cd/m^2^ mean luminance. (B) PVEP amplitudes as a function of spatial frequency for all mice. Linear regressions of PVEP amplitude (blue lines with 95% confidence region) intersect the median noise level (red lines) at different spatial frequencies depending on strain. (C) Group PVEP acuity assessed with categorical logistic method (50% chance of PVEP/noise ratio >1; horizontal arrows represent 50% confidence interval) showing that acuity depends on strain. (D) Mean PVEP acuity (± SEM) in B6 and BALB/c calculated by linear extrapolation of PVEP amplitude to the median noise in the individual mice.

### Visual acuity assessment by optomotor head-tracking response/ reflex (OMR)

Visual acuity was also assessed in B6 and BALB/c mice using the optomotor reflex-based spatial frequency threshold test [8, 16, 17]. In brief, mice were placed on a raised platform at the center of a square box surrounded by four LCD monitors displaying vertical white and black gratings in horizontal motion (clockwise/ counterclockwise direction). A customized program controlled the direction of motion and the stimulus spatial frequency in six steps (0.0275, 0.055, 0.11, 0.22, 0.44 and 0.88 cyc/deg). A camera placed above the mouse monitored head movements, which were viewed on a display by an observer unaware of the direction of the stimulus who scored head-tracking reflex (with head-tracking reflex “1”; without head-tracking reflex “0”). OMRs were scored based on whether the mice tracked the direction of stripe movement with their head. If OMRs were more than 50% correct, then the stimulus spatial frequency was increased stepwise by a factor of 2 until the animal had random OMRs or was unresponsive. Visual acuity was defined as the highest spatial frequency yielding a 50% correct OMR.

### Statistics

All analyses were conducted using JMP Pro 14.2 software (SAS Institute Inc., Cary, NC, USA). PERG and PVEP of amplitudes as a function of spatial frequency were fitted with linear regression, whose intercepts with the median noise level were considered as the corresponding spatial resolution (acuity). PERG and PVEP amplitudes were also expressed as categorical variables for presence or absence of response (SNR >1: YES; SNR ≤ 1: NO) and data submitted to a logistic regression to determine acuity as the highest spatial frequency with at least 50% probability of YES responses. A similar logistic regression was also used to analyze categorical OMR responses.

## Results

### PERG acuity

Examples of PERG waveforms recorded in B6 and BALB/c mice at different spatial frequencies are shown in fig 1A. Fig 1B shows that in both B6 and BALB/c mice, the PERG amplitude at 0.025 -0.05 cyc/deg was on average considerably larger than the median noise level, and then progressively decreased with increasing spatial frequency. For each strain and condition, a linear regression interpolating all data intersected the noise amplitude at a given spatial frequency (PERG acuity). PERG acuities depended on strain and stimulus mean luminance. Average PERG acuities calculated by linearly extrapolating PERG amplitude to the noise level were: 0.53 cyc/deg (B6-700 cd/ m^2^) and 0.22 cyc/deg (B6-70 cd/m^2^) and 0.19 cyc/deg (BALB/c-700 cd/m^2^) and 0.13 cyc/deg (BALB/c-70 cd/m^2^).

PERG acuities were calculated with the same linear regression approach in individual mice (Fig 1C) The average PERG acuities were, B6-700 cd/ m^2^ :0.54 cyc/deg; B6-70 cd/ m^2^ :0.21 cyc/deg; BALB/c-700 cd/ m^2^ :0.18 cyc/deg; BALB/c-70 cd/ m^2^ :0.14 cyc/deg. Statistical analysis showed a strong effect of spatial frequency (P<0.0001), strain (p<0.0001) and luminance (p=0.0007). PERG acuities were also calculated by expressing PERG and noise amplitudes as categorical variables (SNR >1: response; SNR ≤ 1: no response) and using logistic regression to predict the highest spatial frequency with at least 50% probability of acceptable responses (Fig 1D). Logistic regression showed a strong effect of spatial frequency and strain (P<0.001) and a borderline effect of luminance (P=0.05). PERG acuities evaluated with the logistic regression method were, B6-700 cd/ m^2^: 0.54 cyc/deg; B6-70 cd/ m^2^ :0.28 cyc/deg; BALB/c-700 cd/ m^2^ :0.12 cyc/deg; BALB/c-70 cd/ m^2^ :0.06 cyc/deg.

### PVEP acuity

Example of PVEP and noise waveforms obtained from B6 and BALB/c mice are shown in the fig 2A at LED display luminance 700 cd/m^2^. We used three different analytical approaches to validate the PVEP acuities in B6 and BALB/c. The two linear regressions of PVEP amplitudes and median noise extrapolated for group PVEP acuity at 0.55 cyc/deg in B6 and 0.2 cyc/deg in BALB/c mice. (Fig 2B) PVEP signal to median noise ratio was calculated and used ratio 1 as threshold to get the PVEP acuity is 0.55 cyc/deg in B6 and 0.22 cyc/deg in BALB/c by the nominal logistic approach.(Fig 2C) Individual mouse PVEP acuity was also analyzed by linear extrapolation (PVEP amplitudes VS median noises) ; mean PVEP acuity 0.53 cyc/deg in B6 and 0.18 cyc/deg in BALB/c.(Fig 2D)

### OMR acuity

Fig 3 summarizes OMR results. B6 mice exhibited a clear head-tracking movement with the direction of the rotating pattern stimulus as expected, resulting in OMR visual acuity of 0.44 cyc/deg using the categorical logistic approach. BALB /c mice were virtually unresponsive at any spatial frequency. Weak head-tracking responses were occasionally detected but they were random and could not be used to assess visual acuity.

**Fig 3.**
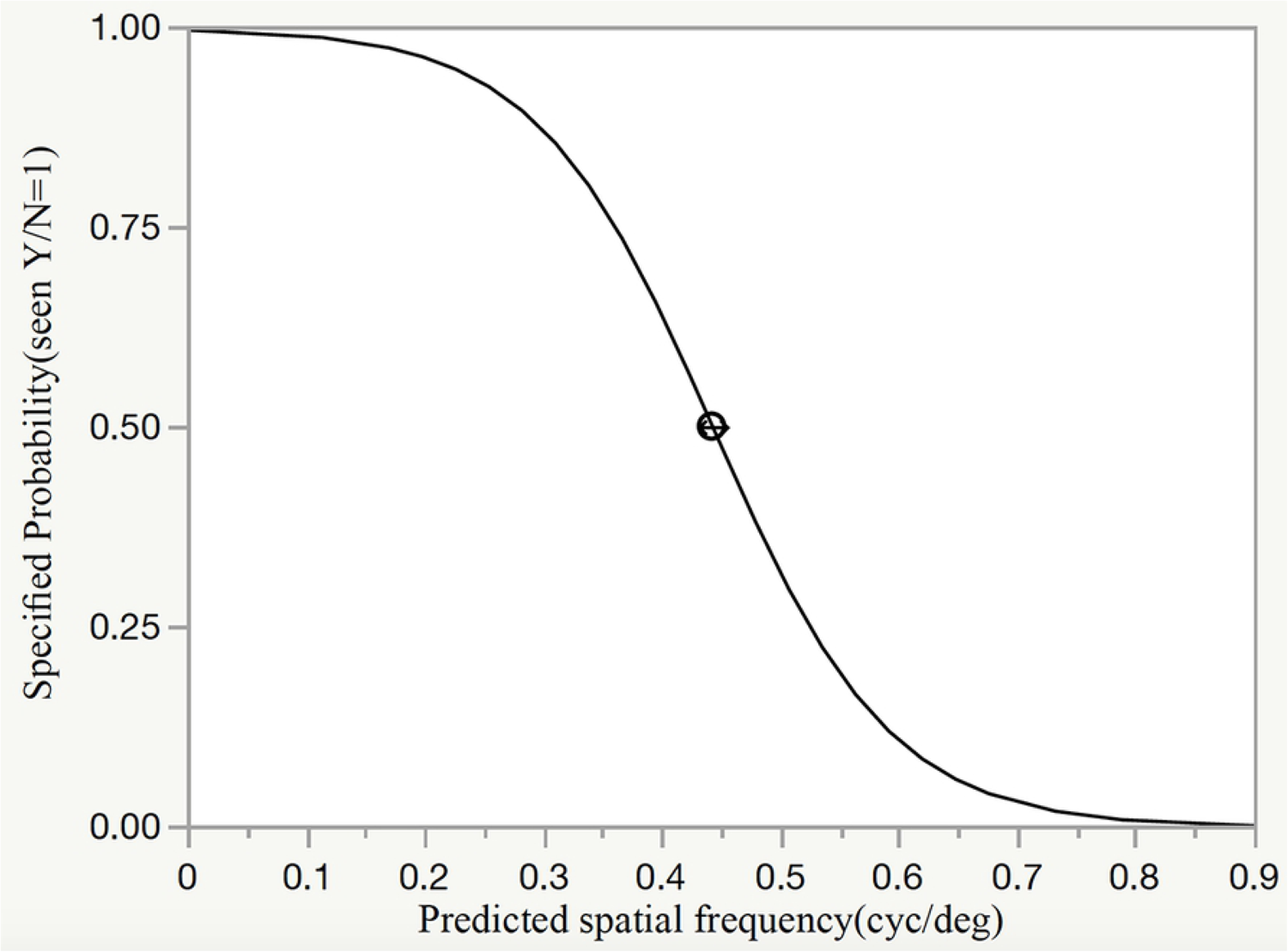
OMR acuity assessed with categorical logistic method in C57BL/6J mice. Logistic regression of BALB/c mice is not shown as they were unresponsive to the OMR stimulus.

## Discussion

Despite the widespread use of inbred albino mice in research involving the visual pathway, few studies have dealt with the actual assessment of visual performance in this strain [5, 7]. Available studies concur that the visual acuity of BALB/c mice is lower than that of pigmented B6 mice, but do not clarify whether the reduced acuity is due to retinal and/or postretinal abnormalities. The present study used combined recording of PERG and PVEP to have an objective assessment of spatial resolution (acuity) at retinal and cortical level. The PERG signal of BALB/c mice, as that of B6 mice, reflects RGC activity as the response is abolished after lesion of the optic nerve that causes RGC degeneration while leaving pre-ganglionic retinal neurons intact [18, 19]. Thus, PERG acuity reflects the spatial resolution of the retinal output. Our results showed that the mean PERG acuity of BALB/c mice was 0.22 cyc/deg and that of B6 mice was 0.55 cyc/deg, calculated with either linear regression of PERG amplitude or logistic regression of categorical SNR>1. By decreasing the stimulus mean luminance from 700 cd/m^2^ to 70 cd/m^2^, mean PERG acuity decreased significantly half in B6 and decreased insignificantly (p=0.76) in BALB/c mice. (Fig 1D) Psychophysical visual acuity reduction with decreasing luminance is a well-known phenomenon in pigmented mammals [20]. However, visual acuities in albinism do not appear to have an obvious luminance-acuity relationship at it has been reported to occur in albinism [21, 22].

In both B6 and BALB/c mice, PERG acuity closely matched PVEP acuity, suggesting that in both strains the visual acuity was primarily determined at retinal level with no further change along the post-retinal visual pathway. Selective loss of spatial resolution in the post-retinal pathway may occur in mutant B6 mice lacking the β2 subunit of the neuronal nicotinic receptor, where PERG acuity is reported to be 0.6 cyc/deg compared to PVEP acuity of 0.3 cyc/deg [12]. In B6 mice, we were able to compare PERG acuity with OMR acuity and found that they were similar. We were unable to elicit a measurable OMR reflex in BALB/c mice, in agreement with other studies [5]. We cannot exclude that a more sophisticated OMR method could have detected small OMR reflexes that allowed assessment of visual acuity [8].

The causes of reduced retinal acuity in BALB/c mice may be associated to the reported reduction of rod density [4, 23], retinal thickness [7, 24], and flash-ERG response [25] that reflects outer retinal activity. Other mechanisms may contribute, as the PERG is a cone-driven response that depends on RGC density as well as intraretinal connectivity [26, 27]. Defects of the post-retinal pathway [7, 28] are less likely to have contributed to reduced retinal acuity.

## Conclusions

The present results show that electrophysiological acuity determined with non-invasive, high-throughput methods that test two eyes simultaneously can be assessed in individual mice in both B6 mice and BALB/c mice. This may be advantageous compared to either operant methods that require training and learning and may not be feasible in some strains or OMR method at which BALB/c mice may be poorly responsive. Also, PERG/PVEP methods may be advantageous compared to both operant and OMR methods as these do not identify the locus of visual defect along the visual pathway.

